# Male-mediated maturation in a wild primate

**DOI:** 10.1101/2020.05.25.114934

**Authors:** Amy Lu, Jacob A. Feder, Noah Snyder-Mackler, Thore J. Bergman, Jacinta C. Beehner

**Author notes:** Correspondence to: Amy Lu, Jacinta C. Beehner.

## Abstract

In humans, a controversial hypothesis suggests that father absence promotes early puberty in daughters. Data from rodents confirm females accelerate maturation with exposure to novel males (“Vandenbergh effect”) and delay it with exposure to male relatives. Here, we report the first case of male-mediated maturation in a wild primate, geladas (*Theropithecus gelada*). Females were more likely to mature after a change in the reproductive male: some matured earlier than expected (Vandenbergh effect) and some later (due to father presence). Novel males stimulated a surge in estrogens for all immature females - even females too young to mature. Although male-mediated puberty accelerated first births, the effect was modest, suggesting that alternative scenarios, such as co-evolution with the Bruce effect (male-mediated fetal loss) may explain this phenomenon.

**One Sentence Summary:** Novel males induce an estrogen surge, male-mediated puberty, and a head-start on reproduction for immature female geladas.

## Main Text

In recent decades, the age at puberty for adolescent girls has been steadily decreasing worldwide (*1*). This trend for earlier puberty has largely been attributed to the growing obesity epidemic (*2*). However, a more controversial hypothesis pervades the human literature – mainly, that adverse childhood experiences, particularly those related to father absence, promote early puberty in adolescents (*3–5*). Although this hypothesis has gained some support (*6–8*), it is difficult in humans to separate the effects of father-absence from the effects of stress-related challenges of a dysfunctional family environment that often (but not always (*9*)) accompany father-absence.

To identify the specific role of the male environment on female maturation, we can look to animal models for answers. More than 50 years ago, John Vandenbergh conducted a series of experiments demonstrating that female mice mature earlier if they are housed with an unrelated adult male (*10*). This “Vandenbergh effect” – the acceleration of maturation by the presence of an unrelated male – has since been found in other rodents, several domestic livestock, and one marsupial (Table 1). The mechanism for the Vandenbergh effect has been well-characterized in rodents, comprising a combination of social and chemical signals (*11–13*). The chemical signals derive from a male’s urine, and is likely to be urinary estradiol (*14, 15*). Both exogenous (*16*) and endogenous (*17*) estradiol are known to promote uterine growth in females. Moreover, in a related phenomenon, females in many rodent taxa experience a delay in maturation when continually exposed to their male kin (mainly fathers and brothers). The male-kin delay in pubertal onset often occurs in species that also exhibit the Vandenbergh effect, suggesting that the two effects may be mechanistically linked (**Table S1**).

Despite debates on whether father-absence accelerates or father-presence delays puberty in humans (*18*), we have surprisingly limited comparative data from primates and none from natural settings. We have two hints from captive primates that the male environment may mediate pubertal changes in females; male turnover was associated with early puberty in two species of galagos (*19*) and in one anecdotal account from hamadryas baboons (*20*). Although the small-bodied, cooperatively breeding callitrichids routinely show socially mediated puberty (*21*), this seems largely driven by the suppressive effects of females, and not males (*22*).

Moreover, despite extensive study of the proximate mechanisms behind male-mediated puberty in rodents, we have no data on the adaptive function of this phenomenon in rodents or in any other taxa. Life history theory suggests that females should optimize the timing of maturation in response to unpredictable social conditions: if only related males are present, females should remain immature and continue to invest energy in growth; however, if an unrelated male arrives, females should mature and shunt energy to reproduction (*23*). Thus, male-mediated maturation may be adaptive because it can accelerate a female’s reproductive career when conditions are optimal. Alternative scenarios are also possible, however. For example, the adaptive benefit of male-mediated maturation may hinge on coinciding the start of a female’s reproductive career with the arrival of a male - for instance, to avoid costs of infanticide in species with high male turnover rates. Testing these hypotheses requires data from longitudinal studies in wild animals, where we can quantify whether accelerated maturation or timing maturation with the arrival of novel males can influence reproductive success.

Here, we used 14 years of demographic and hormone data to test for male-mediated maturation in a wild primate, the gelada (*Theropithecus gelada*). Previously, geladas were shown to exhibit the Bruce effect, a phenomenon where pregnant females spontaneously abort following the arrival of a novel male (*24, 25*). Around the same time, it was shown that the Vandenbergh effect accompanies the Bruce effect in mice and that the same mechanism mediates both effects (*14*); exogenous estradiol from a novel male’s urine causes immature females to accelerate maturation and causes recently inseminated females to abort (*14, 26*). This shared mechanism prompted us to examine the effects of the males on the timing of female puberty in geladas, including acceleration due to novel males and delay due to father-presence. We then tested if estrogens might be associated with male-mediated reproductive maturation, and examined putative benefits of male-mediated reproductive maturation.

To test for male-mediated maturation, we collected demographic and behavioral data from a wild population of geladas living in the Simien Mountains National Park, Ethiopia. Geladas are catarrhine primates that live in polygynous family units (“reproductive units”) comprising one dominant breeding male (“leader male”), 1-12 related, adult females, and their dependent offspring (*27*). Reproductive success for leader males depends on maintaining control over the unit. Threats to leader males come from “bachelor” males residing in all-male groups, who challenge and defeat leader males (“male takeover”), allowing the successful bachelor to gain reproductive access to the unit’s females (*28*). Takeovers are semi-seasonal and each male is replaced approximately every 2.74 years (89 takeovers across 228 unit-years of study) - usually well before his daughters reach maturity around 5 years of age. However, in this population, both male tenure (<1mo - 8.1 years, N=67) and age at maturation (3.5-6.5 years, N=80) are highly variable, creating the potential for overlap. We recorded female maturations based on the first signs of sex skin swelling on a female’s chest (*29*). Such swellings are extremely conspicuous (**Fig. 1a**) and are tightly correlated with changes in fecal estrogens (**Fig. S1**), indicating that they serve as an accurate morphological proxy for reproductive maturation.

**Fig. 1.**
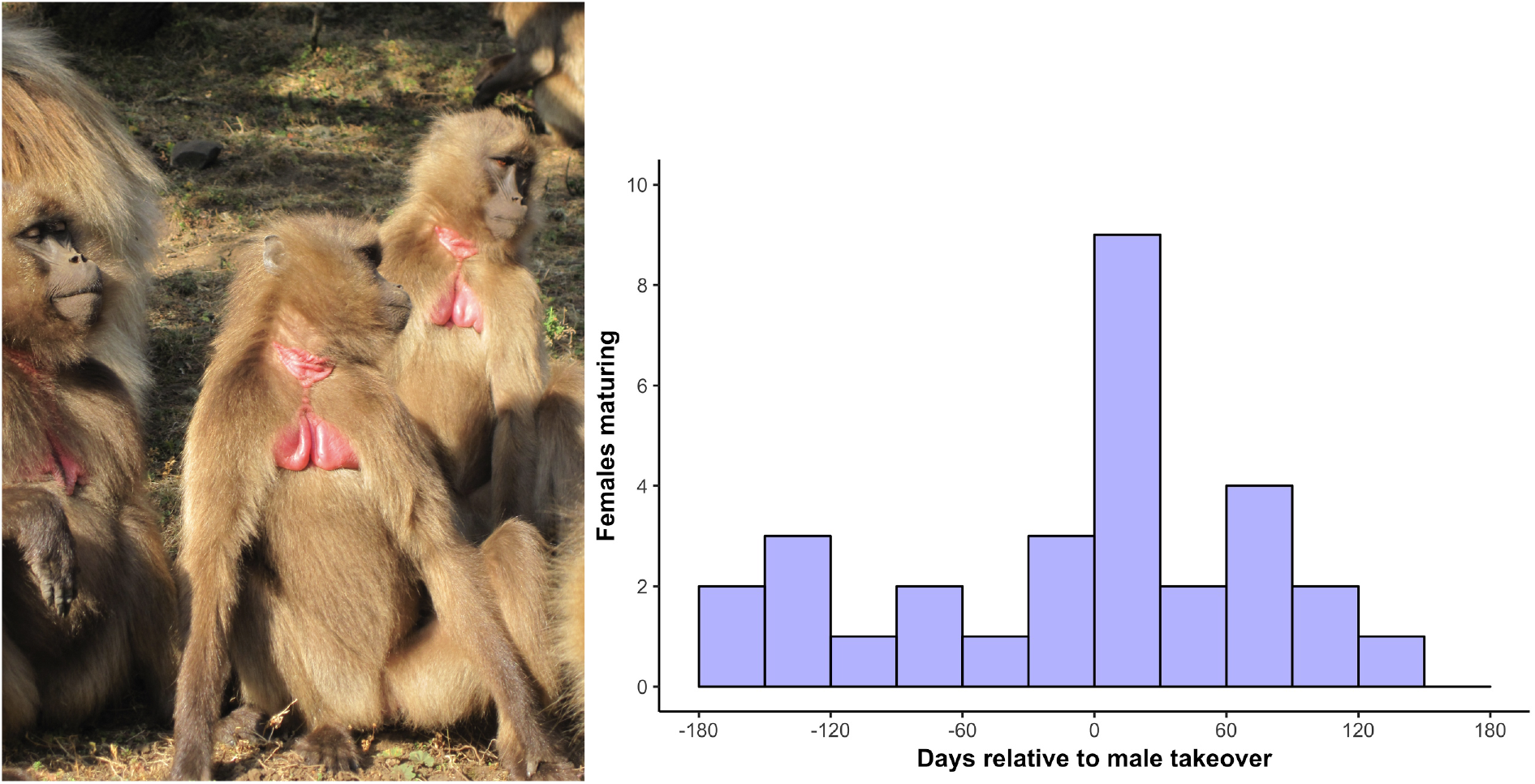
Females were more likely to mature during the three months following a male takeover. The first sexual swelling in geladas for two females (photo by J. Jarvey, with permission)(**1a**). Histogram of 39 female maturations relative to the timing of a male takeover (days, +/- 6 months; **1b**). Histogram data were drawn from a sample of 55 maturations from females of known age and 25 maturations from females with estimated ages. 18.8% of all maturations occurred within the three months following takeovers. Note that maturations that did not occur within +/- 6 months of a takeover are not included in this figure.

First, we analyzed demographic data from 80 immature females across 28 takeovers during a 14-year period (Jan 2006 - Jul 2019) to determine if maturations increased following the arrival of a new male. Females were nearly three times more likely to mature after male takeovers (HR = 2.86, z = 3.25, P = 0.001; **Fig. 1b**). However, females were less likely to mature when their father was still the leader male in their social unit (HR = 0.66, z = −4.49, P < 0.0001; **Fig. S2**). Notably, the hazard ratio of father presence was not consistent over time (*χ*^2^ = 4.66, P = 0.03) indicating that the suppressive effect of fathers decreased with female age. Similarly, older females were more likely to mature following male takeovers than younger females (generalized linear mixed model: β=2.44, z=3.20, P=0.001; **Table S2**). A female experiencing a male takeover at the median age of maturation (4.77 years) would have a 67.5% likelihood of maturing within the next three months; however, by 6 years of age, this effect approaches 100% (**Fig. S3**). Furthermore, while takeovers in general lift the inhibition of maturation in gelada females, the presence of fathers as primary breeding males counters this effect. We found that, on average, male takeovers accelerated maturation by 4.6 months (linear mixed model: β = - 0.383, t =-2.80, P = 0.007; N=80; **Fig. 2, Table S2**), and father-presence delayed maturation by 5.2 months (β = 0.434, t =3.61, P = 0.0005).

**Fig. 2.**
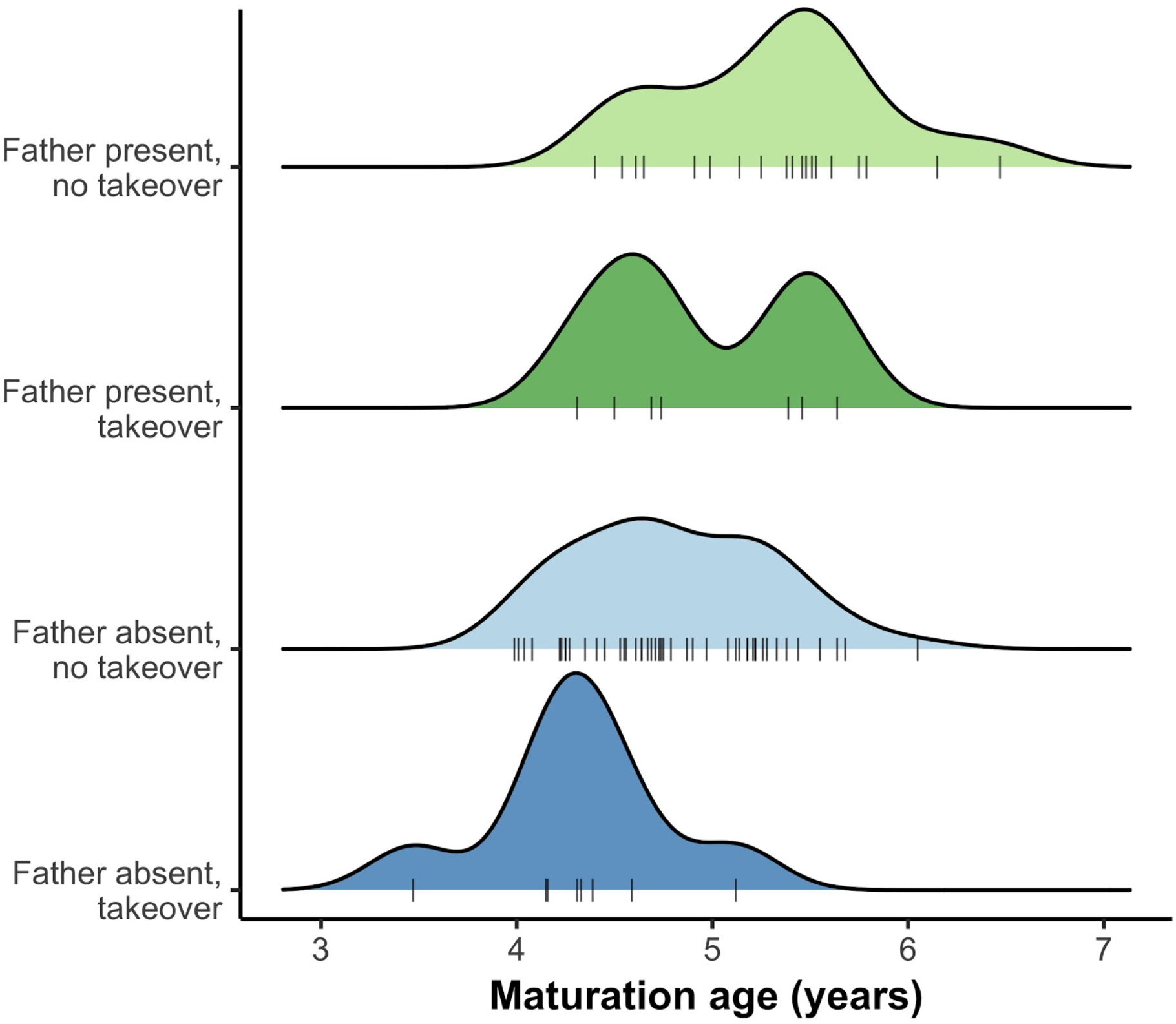
Females matured earlier following takeovers and later if their fathers were still the breeding male. Ridge plot of 80 female maturations ages. Maturations are separated into four groups based on whether females’ fathers were absent (**blue**) or present (**green**) at the minimum age at maturation (41.7 months) and whether females matured following (**light**) or not following (**dark**) a male takeover. The dashed line indicates the median age at maturation for all females (N=80).

Next, we investigated one putative mechanism for the observed male-mediated effects on female maturation. In mice, male urinary estradiol was absorbed by immature females, causing immediate increases in ovarian and uterine weights, which led to maturation (*30*). To examine whether a similar estradiol-based mechanism occurs in geladas, we analyzed hormonal data from 51 juvenile females, including 42 females with known dates of maturations and 9 females that were only observed as juveniles that had yet to mature. Using radioimmunoassay of fecal estrogens (*24*), we tested if the arrival of a new male increased female fecal estrogen metabolites for immature females during the immediate 30 days following the takeover (physiological effects of males are expected to be immediate, preceding subsequent effects on sex skin swellings (**Fig. S1**)).

Outside of takeovers, estrogen metabolites ranged from 0.06-10.31 ng/g; however, following male takeovers, these values more than doubled (pre-takeover mean=1.47; post-takeover mean=3.15 ng/g; **Fig. 3a**). This observation parallels those in mice (*31, 32*): novel males trigger an immediate rise in estrogens for immature females. These effects were not limited to females of maturational age: we found that estrogens increased in immature females of all age classes (β = 0.54, t = 8.54, P < 0.0001; **Fig. 3b, Table S4**) - even females as young as one year old, which is years younger than the earliest age at maturation. Notably, no gelada females under three years of age matured in response to takeovers, indicating that this brief rise in estrogens is not, by itself, sufficient to induce maturation in the youngest females.

**Fig. 3.**
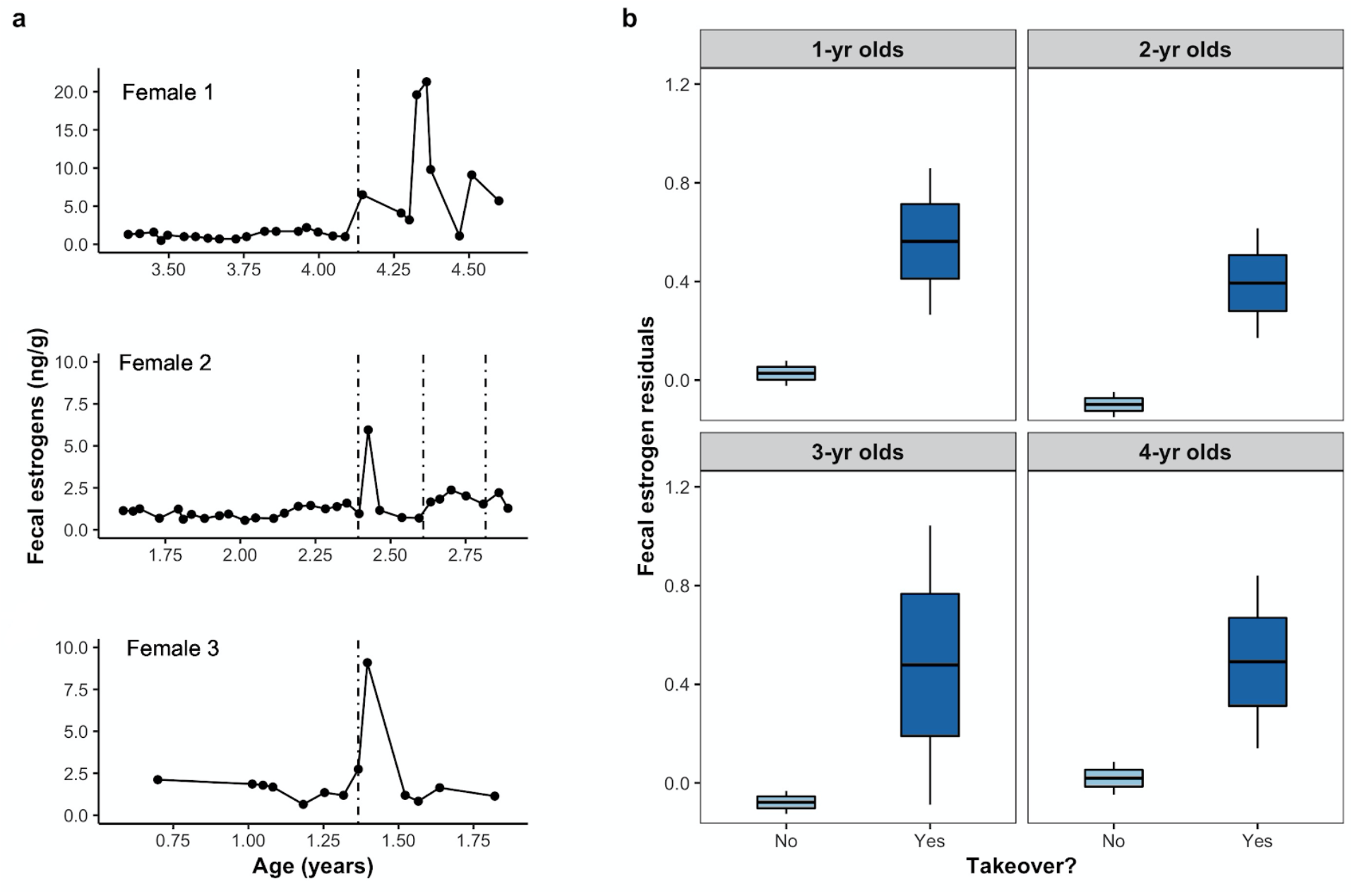
Male takeovers induced increases in estrogen levels in all pre-pubescent females. (A) Fecal estrogen metabolites for three representative adolescent females at different ages (Female 1 is four years old, Female 2 is two and a half years old, and Female 3 is one and a half years old) relative to the timing of male takeovers (indicated by a dashed line). More than one dashed line indicates multiple takeovers (**3a**). Residual fecal estrogen metabolites for females at different ages (1-4 years of age) either outside of (“no”) or within the 30-day window after a takeover (“yes”) (**3b**). Fecal estrogens presented as residuals controlling for female age, cumulative rainfall (previous 90 days), average minimum temperature (previous 30 days), whether the sample was collected within the 100 days before the female’s first swelling, and methodological differences in sample extraction. For (3b), mean=solid line, standard error=box outline, 95% confidence interval=whiskers.

Although the surge in estrogens that accompanies typical mammalian maturation derives from endogenous sources (e.g., ovaries, adrenals), most examples of Vandenbergh-stimulated maturation derive from exogenous sources (e.g., male urinary estrogens). We currently do not know whether the gelada spike in estrogens after takeovers is due to endogenous or exogenous sources. However, we propose that they are endogenous for two reasons: catarrhine primates, like geladas, do not have a functioning vomeronasal organ (*33*), the primary structure that is known to processes chemosignals (*34*), and geladas live in large societies of up to 1200 individuals (*27*), making it difficult for a system of chemical communication to target a specific individual. Catarrhine primates have replaced the chemosensory input used by most other mammals with cognitive processing of environmental cues (*35*). Therefore, it is more likely that such a mechanism derives from socio-cognitive input, followed by downstream neuroendocrine changes in the hypothalamic-pituitary gonadal (HPG) axis that parallel those confirmed in other mammals (e.g., the stimulation of gonadotropin releasing hormone (GnRH) surges) (*36*). Regardless, it appears that females must be physiologically “ready” (i.e., have reached some critical threshold) before estradiol can stimulate reproductive maturation. In prepubertal females, estrogens exert negative feedback on the HPG axis, preventing the GnRH surges necessary for the initiation of puberty (*37*). As females approach the pubertal transition, the inhibitory influence of estrogens on the HPG axis decreases, eventually switching to a stimulatory one that initiates the development of the female reproductive tract, uterine growth (*15*), functional menstrual cycles, sexual behavior, and for some primates, the development of sexual swellings (*38*).

Despite being documented in about two dozen mammalian taxa (**Table S1**), we currently have no evidence that male-mediated maturation is adaptive. There are three possible adaptive benefits. First, the benefits of the Vandenbergh effect may hinge on whether it accelerates the time to first successful reproduction. Indeed, gelada females that matured earlier also had an earlier first birth (β = 0.86, t =7.75, P < 0.0001; N=62, **Fig. S4**). Given their earlier ages at maturation, females that experienced a male-mediated maturation gained a ~4-month head start on reproduction, resulting in a modest 0.14 offspring advantage (based on an interbirth interval of 2.44 years) across a lifetime, all else being equal (e.g., longevity).

Second, this modest head start on reproduction may not be the primary advantage. Male-mediated puberty (regardless of whether it occurs early, on-time, or later) may bestow a timing advantage for female geladas. The arrival of novel males carries two costs for adult females: the Bruce effect for pregnant females (*24*) and infanticide for lactating females (*39*); and selection may therefore favor females that time the start of a reproductive event with the arrival of a new male (*24*). That is, females that time conceptions with male takeovers should have more time to gestate and wean an infant before another male takeover occurs. This timing hypothesis is a bit more difficult to predict for immature female geladas because they exhibit a period of “adolescent sterility” after maturation (1.42 years; N=94 females, SD=0.47 years). Nevertheless, females that initiate this process at the time of male takeovers may have an advantage. We found some support for this hypothesis. Females that matured at the time of a male takeover subsequently “bought” themselves more time to get through this sterile period, conceive, gestate, and wean their first offspring before the next male takeover (linear mixed model: β=359.4; t=2.56, P=0.01; N=63, **Table S5**); such females were takeover free for 1087 days (2.98 years, SE=160 days) following their maturation, while all other females (“control females”) were takeover free for only 727 days (1.99 years, SE=116 days). Thus, on average, females with male-mediated maturation are expected to have a 1-year-old infant at the time of the next takeover (such infants are less susceptible to infanticide (*39*)) while other females are expected to have a newborn infant. Male-mediated maturation may, therefore, help females avoid the Bruce effect and infanticide that accompany male takeovers. To directly test the second of these scenarios, we examined first-infant survival for male-mediated and control maturations. Contrary to expectations, male-mediated puberty did not ensure higher survival for these females’ first infants (survival in relation to all causes of infant mortality: 12/16, 75.0%; survival in relation to takeover-related infant mortality: 2/4, 50%) compared to all other females (all causes: 55/63, 87.3 %, Fisher exact test; P=0.25; takeover-related: 3/8, 37.5%).

Third, we expect that sensitivity to the arrival of novel males is necessary to immediately lift the puberty inhibition caused by father-presence. It is reasonable to assume that the neuroendocrine process that releases delayed-females from puberty inhibition due to father-presence is the same as that releasing all females from inhibition following the arrival of novel males. There is ample support for the selective advantage behind inbreeding avoidance (*40, 41*), with costs as high as high as an 87% reduction in lifetime fitness (*42*). Although we are unable to directly test inbreeding costs in geladas (we only observed one instance of a father mating with his daughter), gelada females lose only 0.85 months until their first birth for every one-month of maturational delay, resulting in 0.15 fewer offspring across a lifetime. The benefits to avoiding inbreeding are therefore likely to outweigh the costs, favoring delayed maturation in the presence of fathers.

Finally, female sensitivity to the arrival of novel males may simply be a by-product of the Bruce effect. The Bruce effect is adaptive in geladas (*24*) and is thought to also be adaptive in other taxa (*43*). Moreover, studies have shown that the Vandenbergh and Bruce effects are mediated by the same mechanism (*14, 26*). Male-mediated maturation may therefore be a consequence of selection for the Bruce-effect. This scenario is particularly likely given that the ability to terminate gestation in response to infanticide risk may be adaptive across the entire lifespan, while maturing in response to a novel male is only adaptive for females’ first-infants. As long as Vandenbergh is not costly, this putative pleiotropy between the two would endure. Thus far, all cases in which the Bruce and Vandenbergh effects have been investigated demonstrate complete overlap between the two phenomena (**Table S1**). However, a proper comparative analysis will depend on filling in missing data and incorporating additional mammalian taxa, particularly to establish evidence of absence of either effect.

In sum, we found that (1) male-mediated maturation occurred in a wild primate, (2) the effect is accompanied (and possibly mediated) by a similar mechanism to rodents, a surge in estrogens; and (3) male-mediated maturation may be adaptive, as it accelerates the time to first reproduction and ensures a female will immediately mature when her father is no longer the breeding male. With respect to synchronizing maturation with the arrival of a new male, females with male-mediated maturation appeared to buy more time to avoid the costs of another male takeover prior to weaning their first offspring. However, this bought time did not translate into higher infant survival, likely because the variance in this extended safety net was high. Remarkably, the Vandenbergh effect co-occurs with the Bruce effect in all cases observed to date. Therefore, even if it bestows limited fitness benefits for immature females, it may simply reflect the sensitivity that females retain to the arrival of new males - a plastic physiological response driven by fitness benefits associated with the Bruce effect. As such, we propose that all male-mediated puberty (not just *accelerated* puberty characterized by previous definitions of the Vandenbergh effect) will accompany the Bruce effect. Further, we propose that the term “Vandenbergh effect” should be broadened to encompass all male-mediated puberty and not just accelerated puberty, as the same process governs puberty onset regardless of whether it is early, on-time, or late.

Taken together, these data indicate that puberty in a non-human primate is highly sensitive to the identity of potential male mates, with novel males accelerating and biological fathers delaying pubertal onset. These data from a catarrhine primate suggest that humans may also have retained sensitivity to individual household males, however, the adaptive purpose of retaining such sensitivity in humans is far less clear. Although accelerated puberty might be generally beneficial, it is difficult to understand how timing puberty to the arrival of new males in human societies might improve fetal or infant survival. Furthermore, given the weak (or nonexistent) evidence for sexually-selected infanticide and the Bruce effect in humans (*44*), it is unlikely to have persisted due to evolutionary pleiotropy.

Finally, we were unable to probe interindividual variation in this sensitivity to males. Beyond a female’s age (which presumably tracks body size and leptin concentrations (*45*)) and father presence, what factors allow one female to lift pubertal inhibition while another does not when faced with a new male? Such analyses (similar to those examining the effects of early life adversity (*46*)) must be restricted to the subset of females that experience a male takeover, comparing those that do mature to those that do not. As with the Bruce effect (*25*), answering this question relies on not only monitoring our gelada population for another 14 years but also documenting the presence and absence of such phenomena across a wide range of mammalian taxa.

## Supporting information

All supplemental materials

## Acknowledgements

We are grateful to the Ethiopian Wildlife and Conservation Authority for granting us permission to conduct this research. We also want to thank the staff and wardens of the Simien Mountains National Park, our Ethiopian staff (Esheti Jejaw, Ambaye Fanta, Setey Girmay, Yeshi Dessie, Tariku W/Aregay, Shifarew Asrat), our research assistants in the field (Clay Wilton, Julie Jarvey, Levi Morris, Tara Regan, Patsy DeLacey, Peter Clark, Evan Sloan, Liz Babbitt, and Maddie Melton), and our laboratory help (Teera Parr, Evelyn Pain).

## Author contributions

**Amy Lu** - Conceptualization, Data curation, Formal analysis, Funding acquisition, Investigation, Methodology, Project administration, Resources, Supervision, Validation, Writing original draft, Writing review and editing.

**Jacob A. Feder** - Formal analysis, Investigation, Methodology, Visualization, Writing original draft, Writing review and editing.

**Noah Snyder-Mackler** - Data curation, Funding acquisition, Project administration, Resources, Supervision, Writing review and editing.

**Thore J. Bergman** - Data curation, Funding acquisition, Project administration, Resources, Supervision, Writing review and editing.

**Jacinta C. Beehner** - Conceptualization, Data curation, Formal analysis, Funding acquisition, Investigation, Methodology, Project administration, Resources, Supervision, Validation, Writing original draft, Writing review and editing.

**Competing interests:** Authors declare no competing interests.

**Data and materials availability:** Data and code are available at https://github.com/GeladaResearchProject/Lu_Vandenbergh_2020.

## Supplementary Materials

Materials and Methods

Figures S1-S4

Table S1-S5

References 46-104

