## Supplementary material for "Male-mediated maturation in a wild primate": All supplemental materials

**This PDF file includes:**

Materials and Methods
Supplementary Text
Figs. S1 to S4
Tables S1 to S5

### Materials and Methods

Hormone collection, extraction, and analysis

Fecal samples (N=1506) were collected within minutes after deposition from positively-identified individuals. Hormones were then extracted from feces using a method described previously (*1*). Specifically, the entire fecal sample was mixed thoroughly with a wooden spatula, and an aliquot of the mixed sample (~ 0.1 g wet feces) was placed in 3 ml of a MeOH/acetone solution (4:1). The solution was immediately homogenized for 1 min using a battery-powered vortexer (BioVortexer, BioSpec Products, Inc., Bartlesville, OK). Approximately 6-8 hours later, 2.5 ml of the fecal homogenate was filtered through a 0.2 μm polytetrafluoroethylene (PTFE) syringeless filter (Whatman, Florham Park, NJ), and the filter was subsequently washed with an additional 1 ml of MeOH/acetone (4:1). We then added 7 ml of distilled water to the filtered homogenate, capped and mixed the solution, and loaded it onto a reverse-phase $C_{18}$ solid-phase extraction cartridge (Sep-Pak Plus, Waters Corporation, Milford, MA). Prior to loading, Sep-Pak cartridges were prepped according to the manufacturer's instructions (with 2 ml MeOH followed by 5 ml filtered water). After the sample was loaded, the cartridge was washed with 1 ml of a sodium azide solution (0.1%). A subset of samples were washed with a 20% MeOH solution (N=371), and this difference was accounted for in all statistical models ("methodological differences"). After loading hormone metabolites on the cartridges, cartridges were allowed to dry at ambient temperature for one week, after which they were stored in Whirl-Pak bags containing ~2 g of silica beads at subzero temperatures (-10˚C) until transported to the University of Michigan for analysis. In the laboratory, steroids were eluted from cartridges with 2.5 ml 100% MeOH and subsequently stored at -20˚C until the time of assay. Dry fecal weights from all samples were obtained to the nearest 0.001g.

All samples were assayed for 17b-estradiol (E2) using a radioimmunoassay (RIA) kit produced by MP Biomedicals. Prior to RIA, all samples were incubated at room temperature for one hour. Then, an aliquot of each sample was evaporated to dryness under nitrogen. Sample aliquots were determined such that hormone metabolite values were within the range of optimal precision of the assay. Kit protocols were followed except that all reagents were halved from the amount suggested by the manufacturer (a common technique employed by researchers measuring fecal steroids to maximize the use of each kit). Internal controls were run in every assay and consisted of a high (binding at 30%) and a low (binding at 70%) "pool" (a composite of many fecal samples). All standards were run in triplicate, all controls and samples were run in duplicate, and mean concentrations are expressed as ng per dry gram of fecal material (ng/g). The MP Biomedicals E2 antibody is known to have minor cross-reactivities with other estrogen metabolites (estrone: 20%; estriol: 1.5%; estradiol-17α: 0.7%). Inter-assay CVs for a low and high sample were 13.4% and 14.2% (N=40) respectively, and intra-assay CVs for the equivalent were 8.74% and 14.73%, respectively (N=10). Assays were conducted in laboratories at both the University of Michigan (N=777 samples) and Stony Brook University (N=729 samples), and this difference was accounted for in all statistical models ("methodological differences").

Statistical analyses

First, to assess the influence of takeovers on the timing of first sexual swelling (N=80), we conducted survival analysis using a time-varying Cox proportional hazards model. For this analysis, females entered the dataset at 3.4 years (40.8 months) of age, just prior to the earliest age at maturation and were modeled on a monthly basis until their maturation. Females' birth dates were either known (N=55) or estimated based on coat color (N=23) or juvenile size (N=2)(*2*). For each female-month, we included takeover status and father presence as binary fixed effects. Based on previous studies (*3*), we assigned "takeover status" as "yes" if a female had been taken over within the previous three months. We assigned "father presence" for each female-month. Because leader males sire between 83-100% of all unit offspring (*4*), a female's father was assigned as the leader male at the time of her conception. In some cases (N=6 females), females experienced multiple takeovers in quick succession, resulting in more than three consecutive months of "yes" for her takeover status. Also, given that the effect of father presence was non-proportional (as determined via Schoenfeld residuals), we used a time transformation on this predictor, assuming linear change in its hazard ratio over female age. To control for environmental conditions, we also included cumulative rainfall (previous 3 months) and average minimum temperature (previous month) as fixed effects, as these respectively are reliable proxies for grass availability and thermoregulatory stress (*5, 6*). To control for repeated measures of individual females, we included a cluster option on female identity. Survival models were constructed using the R package 'survival' (*7*). We also constructed a binomial generalized linear mixed model (GLMM) to identify the effect of age on whether a female matured in the three months following takeover, using all immature female-takeover events as the unit of analysis (N=125 events; N=64 females), and controlling for female ID as a random effect. In a parallel analysis, we determined whether females that matured in response to males (i.e., those that mature within three months of a male takeover) matured earlier than others, using a linear mixed model (LMM). Here, we used two fixed effects: whether fathers were present at the earliest documented age at maturation (3.48 years) and whether the female matured within three months following a takeover. For all mixed models, reproductive unit was included as a random effect.

Second, to determine the immediate physiological effect of male takeovers, we constructed a linear mixed model (LMM), using logged fecal estrogens concentrations (ng/g) as the outcome. Takeover status (previous 30 days), cumulative rainfall, (previous 90 days), average minimum temperature (previous 30 days), subject age, wash step (sodium azide vs. methanol), and laboratory (UM or SBU) were included as fixed effects. Additionally, in order to control for whether the female was in the process of maturing, we included whether the sample was collected within 100 days prior to the female's maturation as a fixed effect. This allowed us to confirm whether all females experienced increased fecal estrogens following male takeovers, regardless of whether they were in the process of maturing. Both individual and unit ID were included as random effects. The residuals from the resulting model were normally distributed. All LMMs were constructed and assessed using the R packages 'lme4' and 'lmerTest' (*8, 9*).

Next, we quantified three potential benefits of male-mediated maturations: First, we examined whether females with male-mediated maturation gave birth earlier on average than females without male-mediated maturation To do this, we determined the relationship between age at maturation and age at first birth using a linear model. We then extrapolated the net effect of male-mediated puberty by multiplying the earlier maturation advantage conferred by male-mediated puberty (4.6 months) by the delay in first birth for every month of delayed maturation (i.e., the slope from the linear model). Second, we investigated two potential consequences of *timing* maturations to the arrival of novel males. Specifically, we constructed an LMM using male-mediated status (Y/N) as a fixed effect to determine whether females with male-mediated maturation (N=11) had longer intervals than control females (N=52) from the day of maturation until the next male takeover, using unit as a random effect. We excluded male takeovers within 60 days of maturation, since no females had conceived within such a short interval following maturation, and therefore these females would not incur reproductive costs. Then, we assessed firstborn survival using a Fisher exact test, comparing females with male-mediated maturation to control females, excluding all cases where the fate of firstborn offspring was unknown (N=79). Among the offspring that died before weaning, we also examined whether these infant deaths occurred within the six months following takeover, which would suggest infanticide (*10*).
For all survival models and linear models, variance inflation factors (VIFs) were less than 2.0. All statistical analyses were performed in R v.3.6.0 (*11*).

**Figures S1-S4:**

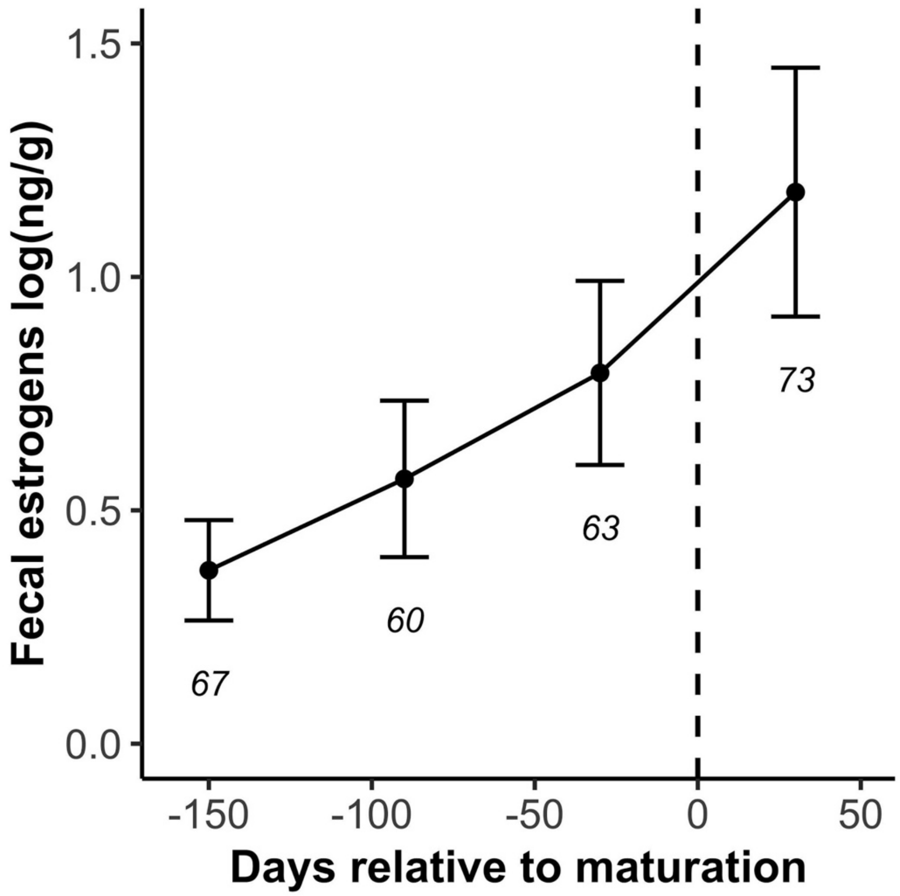


**Fig. S1.** Fecal estrogen concentrations (ng/g) relative to the days surrounding the first onset of sexual swelling for this population of geladas. Hormone sample size presented below error bars.

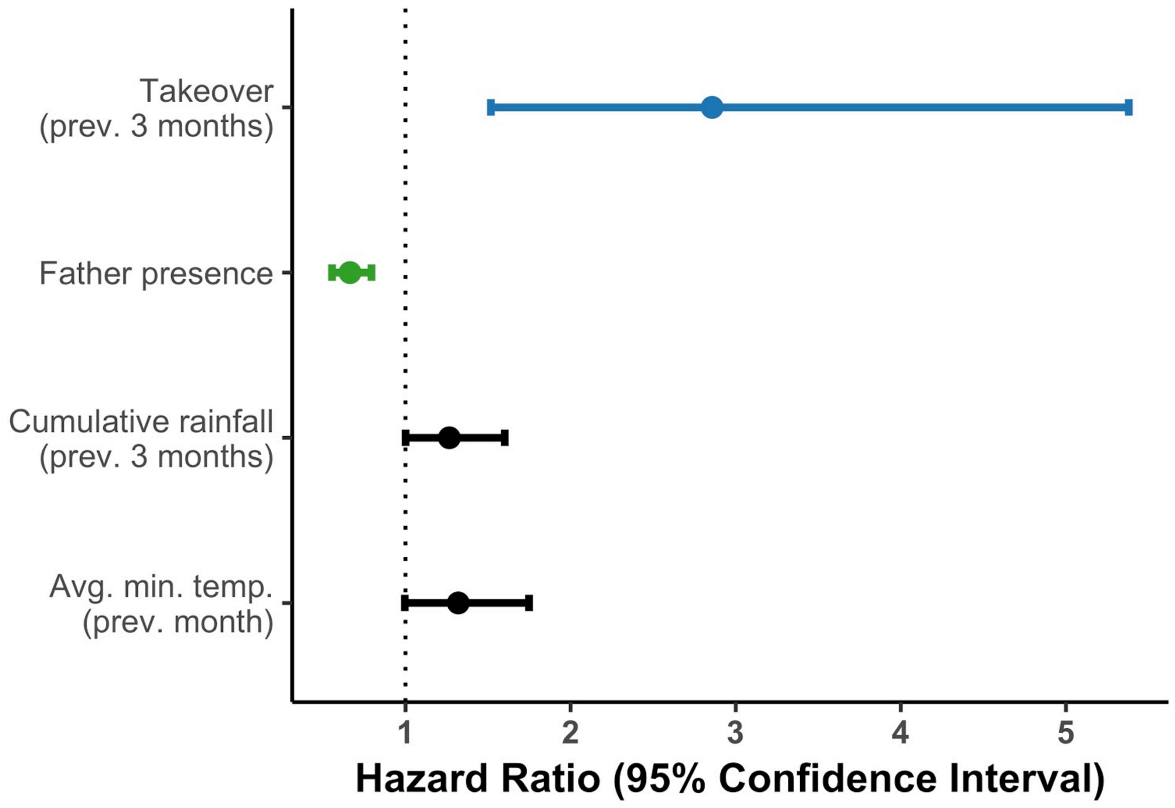


**Fig. S2. Females are more likely to mature in the three months following male takeovers and less likely to mature when fathers are still present in the social unit.** Forest plot of survival model-predicted hazard ratio and 95% confidence intervals. Ecological control variables showed slight trends: females tended to mature at times when there was more cumulative rainfall and higher minimum temperatures (i.e., when grass is highly available and when cold stress is minimal).

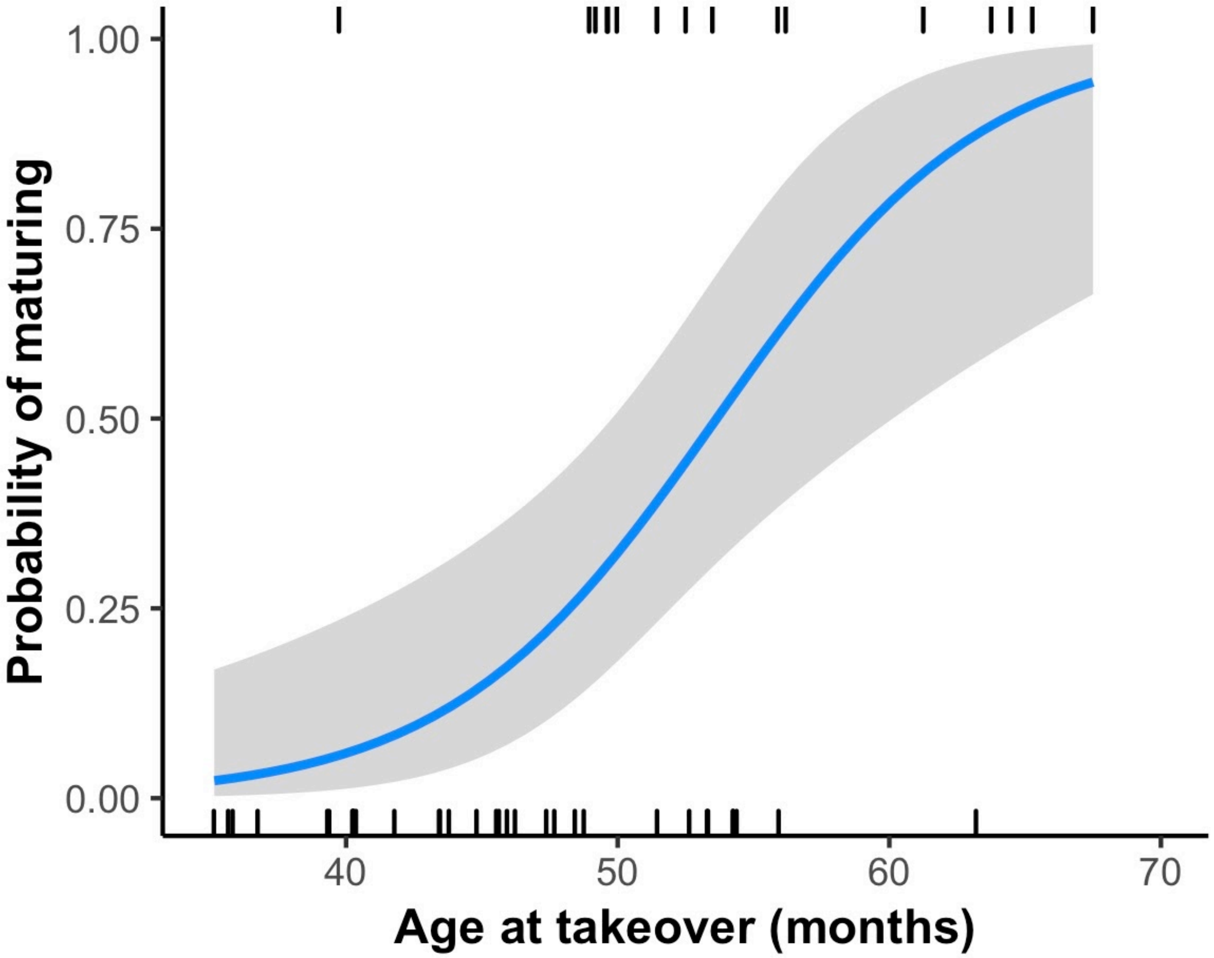


**Fig. S3. Females are more likely to mature following male takeovers as they age** (N=64).

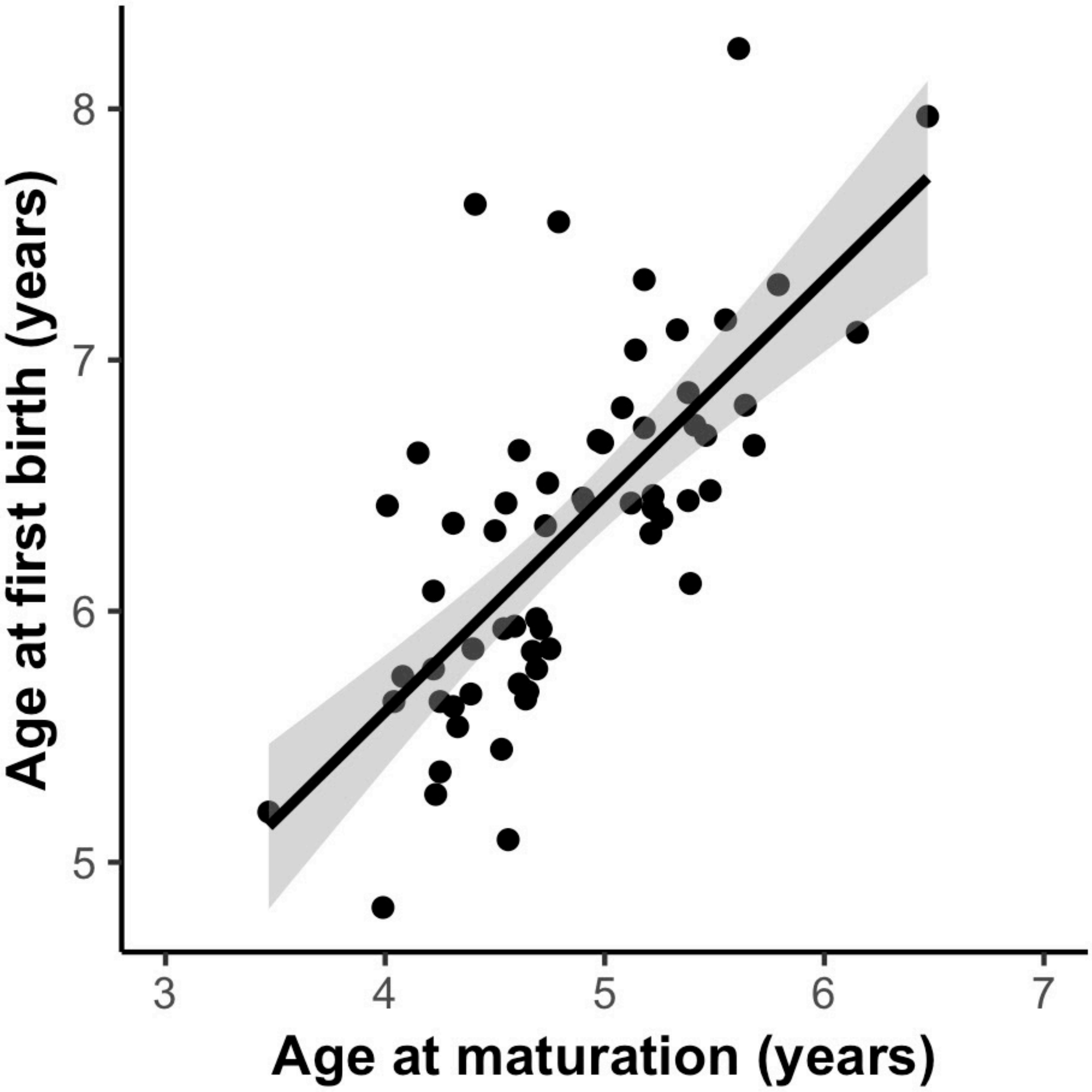


**Fig. S4. Female geladas that mature at earlier ages also have their first birth earlier** (N=62).

### Tables S1-S5:

**Table S1.** All mammalian taxa with the Bruce effect, the Vandenbergh effect (male-accelerated), and male-delayed puberty; co-occurrence of Bruce and Vandenbergh effects are outlined.

| Common name | Latin name | Bruce effect | Ref* | Novel male accelerated puberty | Ref* | Male kin delayed puberty | Ref* |
|---|---|---|---|---|---|---|---|
| **Rodents** | | | | | | | |
| House mouse | *Mus musculus* | yes | (12, 13) | yes | (14) | - | - |
| Hopping mouse | *Notomys alexis* | - | - | yes | (15) | - | - |
| Norway rat | *Rattus norvegicus* | yes | (16) | yes | (17) | - | - |
| Southern African Vlei rat | *Otomys irroratus* | yes | (18) | - | | - | - |
| Deer mouse | *Peromyscus maniclatus* | yes | (19) | yes | (20) | - | - |
| Mongolian gerbil | *Meriones unguiculatus* | yes | (21) | - | | - | - |
| Prairie vole | *Microtus ochragaster* | yes | (22) | yes | (23) | yes | (23) |
| Pine / Woodland vole | *Microtus pinetorum* | yes | (24, 25) | yes | (26) | yes | (27) |
| Meadow vole | *Microtus pennsylvanicus* | yes | (22) | yes | (28) | no | (29) |
| California vole | *Microtus californicus* | yes | (30) | yes | (29) | yes | (29) |
| Montane vole | *Microtus montanus* | yes | (25) | - | - | - | - |
| Field vole | *Microtus agrestis* | yes | (31) | - | - | - | - |
| Tundra / Root vole | *Microtus oeconomus* | yes | (32) | - | - | - | - |
| Sagebrush vole | *Lemmiscus curtatus* | no | (25) | - | - | - | - |
| Brandt's vole | *Lasiopodomys brandti* | yes | (33) | - | - | - | - |
| Southern red-backed vole | *Myodes gapperi* | yes | (34) | - | - | - | - |
| Bank vole | *Myodes glareolus* | yes | (35) | - | - | - | - |
| Collared lemming | *Dicrostonyx groenlandicus* | yes | (36) | yes | (37) | - | - |

| | | | | | | | |
|---|---|---|---|---|---|---|---|
| Norwegian lemming | *Lemmus lemmus* | yes | (38) | - | | - | - |
| Djungarian hamster | *Phodopus sungorus* | - | - | yes | (39) | - | - |
| Syrian hamster | *Mesocricetus auratus* | no | (40) | - | - | - | - |
| Golden hamster | *Mesocricetus auratus* | no | (41) | - | - | - | - |
| Guinea pig | *Galea musteloides* | - | - | yes | (42) | - | - |
| Cavy | *Cavia aperea* | - | - | yes | (43) | - | - |
| Prairie dog | *Cynomys ludovicianus* | - | - | yes | (44) | yes | (44) |
| Cape ground squirrel | *Xerus inauris* | - | - | yes | (45) | - | - |
| Alpine marmot | *Marmota marmota* | yes | (46) | - | - | - | - |
| **Marsupials** | | | | | | | |
| Gray short-tailed opossum | *Monodelphis domestica* | - | - | yes | (47, 48) | - | - |
| **Ungulates** | | | | | | | |
| Pig | *Sus scrofa* | - | - | yes | (49, 50) | - | - |
| Cattle | *Bos taurus* | - | - | yes | (51) | - | - |
| Sheep | *Ovis aries* | yes | (52) | yes | (53) | - | - |
| Creole goat | *Capra aegagrus hirus* | - | - | yes | (54) | - | - |
| Alpine goat | *Capra aegagrus hirus* | - | - | yes | (55) | - | - |
| Domestic horse | *Equus ferus caballus* | yes | (56) | - | - | - | - |
| Domestic (feral) horse | *Equus ferus caballus* | possibly | (57) | - | - | - | - |
| Plains zebra | *Equus berchelli* | possibly | (58) | - | - | - | - |
| Cape mountain zebra | *Equus zebra zebra* | yes | (59) | - | - | - | - |
| **Carnivores** | | | | | | | |
| Domestic dog | *Canis lupus familiaris* | yes | (60) | - | - | - | - |

| | | | | | | | |
|---|---|---|---|---|---|---|---|
| Lion | *Panthera leo* | yes | (*61*) | - | - | - | - |
| **Primates** | | | | | | | |
| Human | *Homo sapiens* | - | - | possibly | (*62*) | possibly | (*62*) |
| Northern lesser galago | *Galago senegalensis* | - | - | yes | (*63*) | yes | (*63*) |
| Garnett's bushbaby | *Otolemur garnettii* | - | - | yes | (*63*) | yes | (*63*) |
| Crested macaques | *Macaca nigra* | no | (*64*) | - | - | - | - |
| Hamadryas baboon | *Papio hamadryas* | possibly | (*65*) | yes | (*65*) | - | - |
| Gelada | *Theropithecus gelada* | yes | (*1*) | yes | *This study* | yes | *This study* |

* References include first reports only

**Table S2.** Results of a generalized linear mixed model examining the influence of age on whether a female matured within three months after experiencing a takeover.

| Fixed effect | Estimate | Std. Error | t-value | P-value |
|---|---|---|---|---|
| (Intercept) | -10.9 | 3.28 | -3.31 | 0.0009 |
| Age at takeover | 2.43 | 0.76 | 3.20 | 0.0014 |

**Table S3.** Results of a linear mixed model examining the influence of takeover status at maturation (i.e., takeover within 3 previous months, Y/N) and father presence when approaching maturation age (41.7 months) on the age at maturation.

| Fixed effect | Estimate | Std. Error | t-value | P-value |
|---|---|---|---|---|
| (Intercept) | 4.86 | 0.08 | 55.2 | <0.0001 |
| Takeover (prev. 3 months) | -0.38 | 0.14 | -2.80 | 0.0066 |
| Father presence | 0.43 | 0.12 | 3.61 | 0.0005 |

**Table S4.** Results of a linear mixed model examining the influence of social, ecological, and methodological predictors on logged fecal estrogens.

| **Fixed effect** | **Estimate** | **Std. Error** | **t-value** | **P-value** |
|---|---|---|---|---|
| (Intercept) | -0.085 | 0.077 | -1.11 | 0.269 |
| Takeover (prev. 30 days) | 0.542 | 0.063 | 8.54 | < 0.0001 |
| First sexual swelling (following 100 days) | 0.449 | 0.061 | 7.32 | <0.0001 |
| Female age | 0.026 | 0.017 | 1.54 | 0.125 |
| Cumulative rainfall (prev. 90 days) | -0.098 | 0.017 | -5.93 | <0.0001 |
| Average min. temp. (prev. 30 days) | -0.037 | 0.014 | -2.68 | 0.007 |
| Wash step* | 0.221 | 0.044 | 5.06 | <0.0001 |
| Laboratory†+ | 0.094 | 0.038 | 2.45 | 0.014 |

*Reference category: MeOH wash; †+Reference category: Stony Brook University

**Table S5.** Results of a linear mixed model examining the influence of male-mediated puberty on the length of the interval between maturation and the first subsequent takeover.

| Fixed effect | Estimate | Std. Error | t-value | P-value |
|---|---|---|---|---|
| (Intercept) | 727.3 | 114.9 | 6.33 | <0.0001 |
| Male-mediated (Y/N) | 359.4 | 140.3 | 2.56 | 0.013 |
